# Fine-mapping of genetic loci driving spontaneous clearance of hepatitis C virus infection

**DOI:** 10.1101/128975

**Authors:** Hailiang Huang, Priya Duggal, Chloe L. Thio, Rachel Latanich, James J. Goedert, Alessandra Mangia, Andrea L. Cox, Gregory D. Kirk, Shruti Mehta, Joel N. Blankson, Jasneet Aneja, Laurent Alric, Sharyne M. Donfield, Matthew E. Cramp, Salim I. Khakoo, Leslie H. Tobler, Michael Busch, Graeme J. Alexander, Hugo R. Rosen, Brian R. Edlin, Georg M. Lauer, David L. Thomas, Mark J. Daly, Raymond T. Chung, Arthur Y. Kim

## Abstract

Approximately three quarters of acute HCV infections evolve to a chronic state, while one quarter are spontaneously cleared. Genetic predispositions strongly contribute to the development of chronicity. We have conducted a genome-wide association study to identify genomic variants underlying HCV spontaneous clearance using Immunochip in European and African ancestries. We confirmed two previously reported significant associations, in the *IL28B/IFNL4*^1,2^ and MHC regions, with spontaneous clearance in the European population. We further fine-mapped the MHC association to a region of about 50 kilo base pairs, down from 1 mega base pairs in the previous study. Additional analyses suggested that the association in the MHC locus might be significantly stronger for virus subtype 1a than 1b, suggesting that viral subtype may have influenced the genetic mechanism underlying the clearance of HCV.

---

The development of chronic viral infection represents a failure to mount an adequate innate and/or adaptive response to a specific pathogen. Infection with hepatitis C virus (HCV) in humans represents a paradigm of a dichotomous outcome of infection, as approximately three quarters of acute HCV infections evolve to a chronic state, but one quarter are spontaneously cleared^3^. As such, it is likely that genetic predispositions, especially at loci that regulate the innate and/or adaptive immune response, strongly contribute to the development of chronicity. A prior genome wide association study (GWAS) conducted by our consortium demonstrated striking associations of spontaneous resolution of HCV with polymorphisms near the IFN-L3 locus (*IL28B*) and in the HLA class II locus (Duggal *et al.*^4^).

The associations identified in Duggal *et al.* span large genomic regions, specifically 55,000 base pairs for the Il28B locus and > 1 mega base pairs for the HLA class II locus. Recently advances in genomic technologies allowed a more precise characterization of genetic associations and facilitated resolving these associations to much smaller genomic regions. Firstly, Immunochip^5^, a customized array platform with deeper coverage of loci enriched in autoimmune diseases, provides coverage of additional genomic variants for an opportunity to explore with greater precision the contribution of these loci to the clearance of viral infection. Secondly, additional coverage of the MHC region can be gained using an imputation algorithm that takes into account the long range linkage-disequilibrium in MHC, and a large customized reference panel with improved coverage of the MHC region^6^. Thirdly, fine-mapping algorithms^7,8^, designed with the goal to resolve known genetic associations to smaller sets of variants, can be used with the high density genomic data to further improve the precision of the genetic associations.

We therefore conducted an analysis of a large pool of spontaneous resolvers and chronic patients of HCV using the Immunochip platform, the SNP2HLA algorithm with the T1DGC MHC imputation reference panel^6^, and a recently-developed fine-mapping algorithm^8,9^ to (1) more precisely define the susceptible variant within the known associated loci; and (2) identify additional loci associated with clearance. Similar successes have been achieved in other diseases such as inflammatory bowel disease^8,10^. Additionally, we explored the hypothesis that there are shared mechanisms that define a “brisk” immunity able to confer both susceptibility to autoimmune disease and improved control of pathogens. We also examined the influence of region (North America versus European) upon associations with HLA within the European ancestry, as previous studies have shown variability of results, especially for the class I locus^11-14^.

## Results

The final dataset after QC has 166,537 variants for 527 cases/828 controls of European ancestry; and 171,161 variants for 75 cases/171 controls of African ancestry (Table 1). For each ancestry, we performed logistic regression under the additive model using the first two principal components as covariates. The QQ plot (Figure 1, using common variants with >2% minor allele frequency) and the genomic control (GC) factors (0.98 for the European ancestry and 0.92 for the African ancestry using designated null variants) indicate the effective control of the population stratification.

**Table 1.** Variants and samples in this study.

|  | European ancestry | African ancestry |
| --- | --- | --- |
| **Variants** |  |  |
| Initial | 191,357 | 191,357 |
| HWE (p-value <1E-6) | -394 | -300 |
| Missingness (5%) | -24,426 | -19,896 |
| After QC | 166,537 | 171,161 |
| **Samples** |  |  |
| Initial | 1,416 | 227 |
| Missingness (5%) | -24 | -2 |
| Heterozygosity | -9 | -1 |
| Duplicated | -28 | -6 |
| After QC | 527 cases/828 controls | 75 cases/171 controls |

Negative numbers indicate numbers of variants or samples removed.

**Figure 1.**
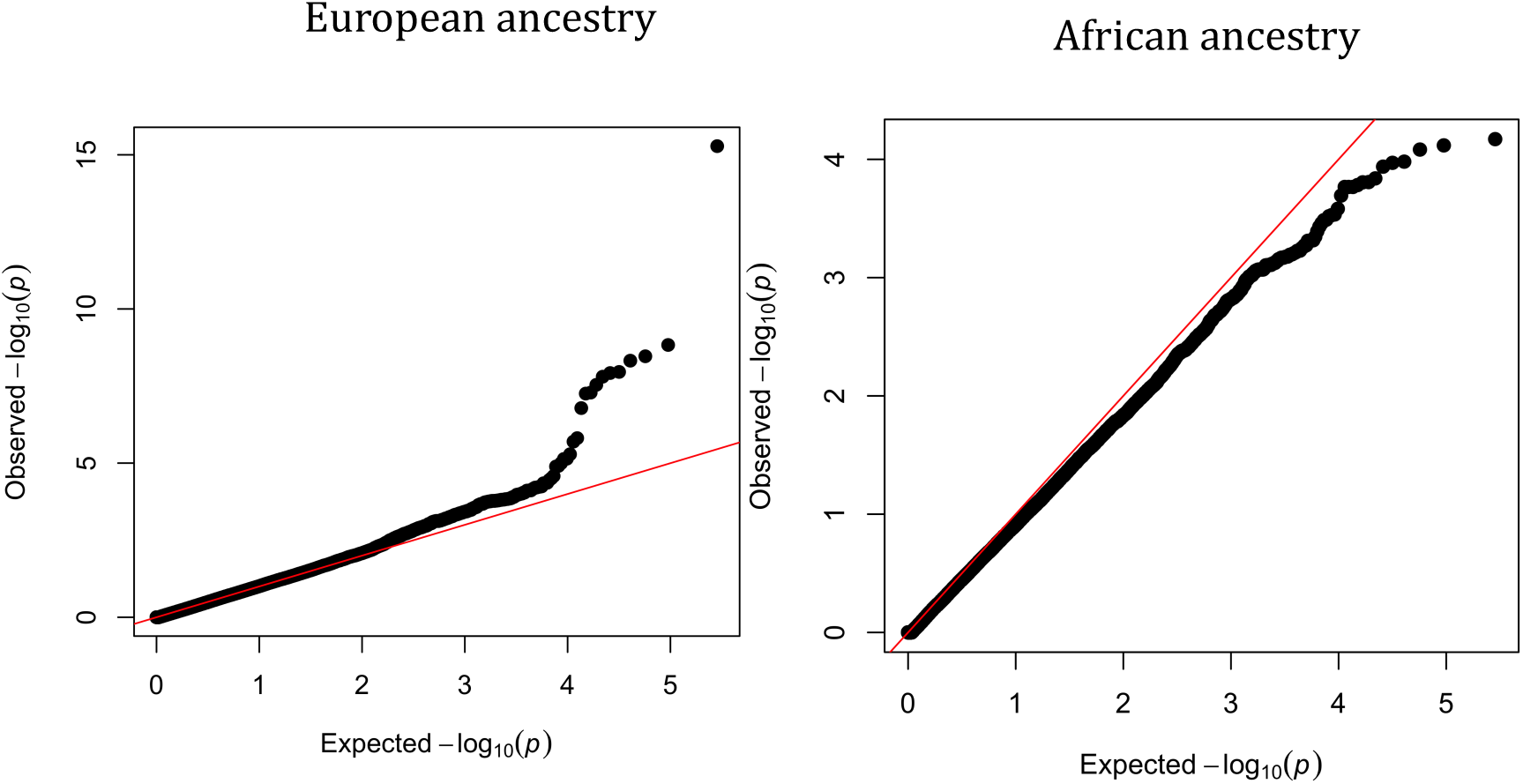
**QQ plot for cohorts of European and African ancestries.** The red line indicates the null distribution. Only variants with minor allele frequency >2% were used in this figure.

For European samples, we identified 8 genome-wide significant variants (p-value < 5E-8) in two loci (Figure 2 and Table 2). The variant on chromosome 19, rs8099917, shows the strongest association with spontaneous clearance (p-value=5E-16). Patients carrying the minor allele, G, are on average 2.5x (odds ratio=0.39) less likely to spontaneously clear the virus compared to those with two copies of the C allele. This variant is roughly 7,000 base pairs upstream of the *IL28B* gene, and has been previously reported to be associated with HCV spontaneous clearance^4^ and the response to chronic HCV therapy in Asian populations^15^. Previous studies have also shown an association between *IL28B* and interferon-based clearance of HCV^16^. Because the *IL28B/IFLN4* region was not designed as a high-density locus in Immunochip, we could not test other variants in this region for their association with HCV spontaneous clearance, and was unable to provide a better resolution in this locus.

**Table 2.**
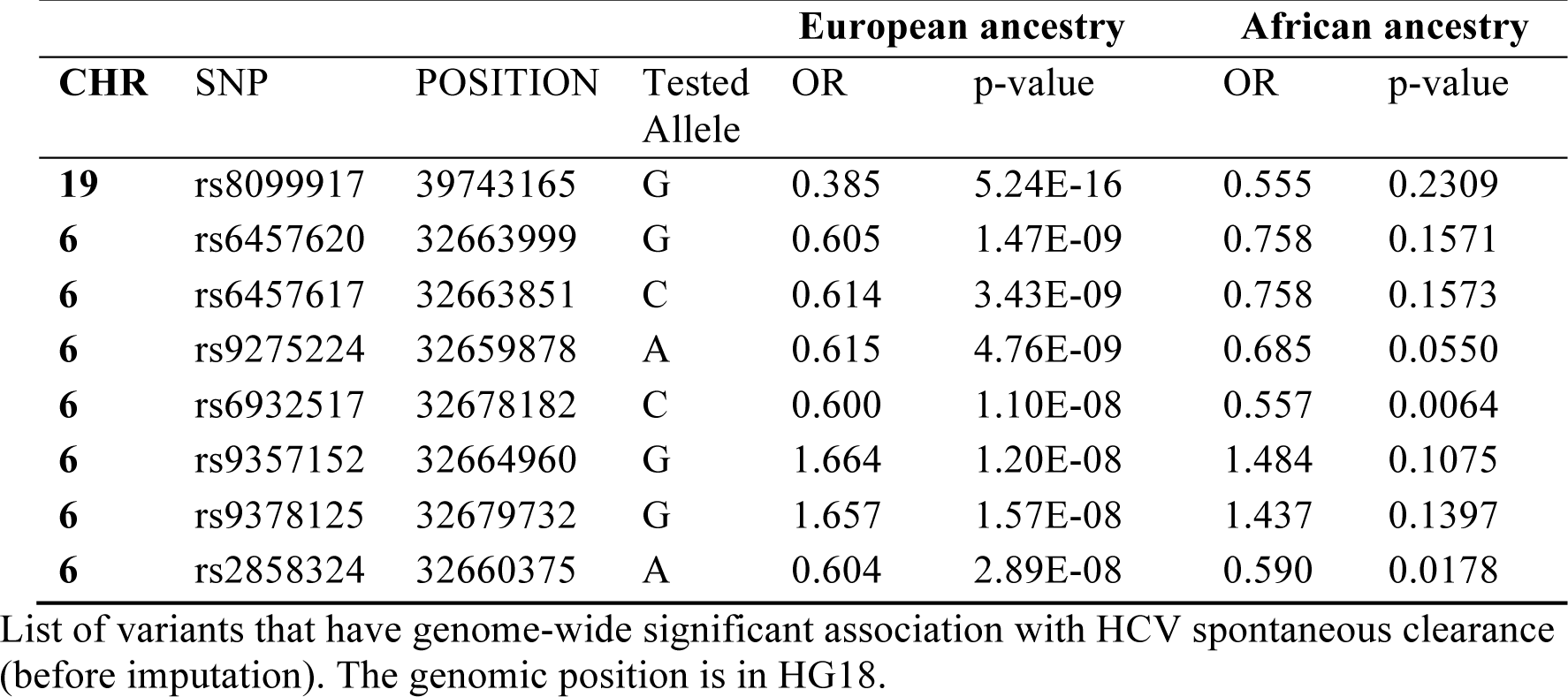
Genome-wide significant associations.

| CHR | SNP | POSITION | Tested Allele | European ancestry |  | African ancestry |  |
| --- | --- | --- | --- | --- | --- | --- | --- |
|  |  |  |  | OR | p-value | OR | p-value |
| **19** | rs8099917 | 39743165 | G | 0.385 | 5.24E-16 | 0.555 | 0.2309 |
| **6** | rs6457620 | 32663999 | G | 0.605 | 1.47E-09 | 0.758 | 0.1571 |
| **6** | rs6457617 | 32663851 | C | 0.614 | 3.43E-09 | 0.758 | 0.1573 |
| **6** | rs9275224 | 32659878 | A | 0.615 | 4.76E-09 | 0.685 | 0.0550 |
| **6** | rs6932517 | 32678182 | C | 0.600 | 1.10E-08 | 0.557 | 0.0064 |
| **6** | rs9357152 | 32664960 | G | 1.664 | 1.20E-08 | 1.484 | 0.1075 |
| **6** | rs9378125 | 32679732 | G | 1.657 | 1.57E-08 | 1.437 | 0.1397 |
| **6** | rs2858324 | 32660375 | A | 0.604 | 2.89E-08 | 0.590 | 0.0178 |

List of variants that have genome-wide significant association with HCV spontaneous clearance (before imputation). The genomic position is in HG18.

**Figure 2.**
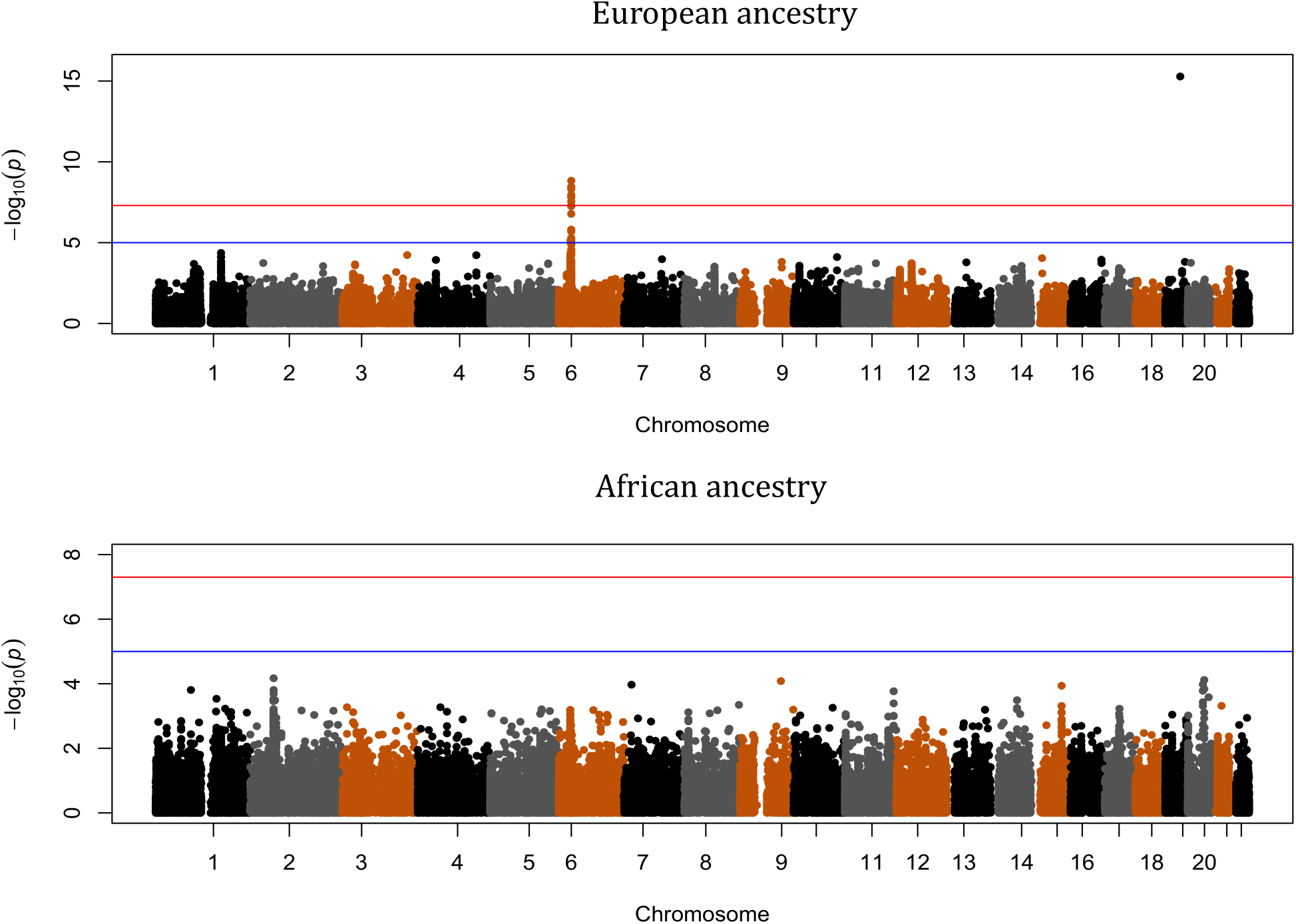
**Manhattan plot for cohorts of European and African ancestries.** The blue horizontal line indicates the suggestive significance threshold, and the red horizontal line indicates the genome-wide significance threshold.

The other genome-wide significant locus for the European samples is the major histocompatibility complex (MHC) locus. Genome-wide significant variants in this region are reported in Table 2 (before imputation). We used SNP2HLA^6^ and a customized reference panel from a T1D study to impute missing variants, HLA alleles and amino acid residues for this region. We identified 12 SNPs and 5 amino acids that are genome-wide significant (Figure 3 and Table 3, boldfaced). No secondary signal in this region exceeded the suggestive significance threshold (1x10^-5^) after conditioning on the primary signal. Therefore, all variants reported in Table 3 account for the same association signal. Using a fine-mapping algorithm described in another study^9^, we constructed the 99% credible set, which is a set of variants that has 99% probability of having the causal variant in this locus (Table 3, full). Comparing with the previous study^4^ which identified this association to a region of more than 1 mega base pairs, we mapped this association to a much smaller region of 50,562 base pairs.

**Table 3.** Associations in the 99% credible set in the MHC region.

| CHR | SNP | POSITION | Tested Allele | OR | P | Probability |
| --- | --- | --- | --- | --- | --- | --- |
| 6 | **rs6457617** | 32771829 | C | 0.6172 | 8.06E-09 | 0.1197 |
| 6 | **rs6457620** | 32771977 | G | 0.6172 | 8.06E-09 | 0.1197 |
| 6 | **rs9378125** | 32787710 | G | 1.664 | 1.34E-08 | 0.0732 |
| 6 | **rs5000632** | 32771666 | G | 1.665 | 1.35E-08 | 0.0727 |
| 6 | **rs9357152** | 32772938 | G | 1.665 | 1.35E-08 | 0.0727 |
| 6 | **rs9394113** | 32773145 | G | 1.665 | 1.35E-08 | 0.0727 |
| 6 | **rs9275224** | 32767856 | A | 0.6229 | 1.60E-08 | 0.0616 |
| 6 | **AA_DQB1_167_32737787_R** | 32737787 | A | 1.717 | 2.05E-08 | 0.0483 |
| 6 | **AA_DQB1_13_32740798_G** | 32740798 | A | 1.708 | 3.03E-08 | 0.0331 |
| 6 | **rs9275516** | 32782621 | A | 0.6079 | 3.60E-08 | 0.0280 |
| 6 | **SNP_DQB1_32737787** | 32737787 | T | 1.7 | 3.79E-08 | 0.0266 |
| 6 | **SNP_DQB1_32740798** | 32740798 | G | 1.7 | 3.79E-08 | 0.0266 |
| 6 | **SNP_DQB1_32740759_T** | 32740759 | P | 1.7 | 3.79E-08 | 0.0266 |
| 6 | **SNP_DQB1_32740760_A** | 32740760 | P | 1.7 | 3.79E-08 | 0.0266 |
| 6 | **AA_DQB1_13_32740798_A** | 32740798 | P | 1.7 | 3.79E-08 | 0.0266 |
| 6 | **AA_DQB1_26_32740759_Y** | 32740759 | P | 1.7 | 3.79E-08 | 0.0266 |
| 6 | **AA_DQB1_167_32737787_H** | 32737787 | P | 1.7 | 3.79E-08 | 0.0266 |
| 6 | **rs6932517** | 32786160 | C | 0.6123 | 5.38E-08 | 0.0190 |
| 6 | **SNP_DQB1_32737837** | 32737837 | A | 0.627 | 8.09E-08 | 0.0128 |
| 6 | **SNP_DQB1_32737148** | 32737148 | G | 1.68 | 9.42E-08 | 0.0110 |
| 6 | **AA_DQB1_45_32740702** | 32740702 | E | 1.68 | 9.42E-08 | 0.0110 |
| 6 | **SNP_DQB1_32740702** | 32740702 | T | 1.68 | 9.42E-08 | 0.0110 |
| 6 | **SNP_DQB1_32742309_A** | 32742309 | P | 1.68 | 9.42E-08 | 0.0110 |
| 6 | **HLA_DQB1_0301** | 32739039 | P | 1.675 | 1.14E-07 | 0.0092 |
| 6 | **rs9469220** | 32766288 | G | 0.6393 | 2.24E-07 | 0.0048 |
| 6 | **rs2856717** | 32778286 | T | 0.6255 | 2.38E-07 | 0.0045 |
| 6 | **AA_DQB1_26_32740759_L** | 32740759 | A | 1.534 | 5.32E-07 | 0.0021 |
| 6 | **SNP_DQB1_32740759_A** | 32740759 | A | 1.534 | 5.32E-07 | 0.0021 |
| 6 | **SNP_DQB1_32740760_G** | 32740760 | A | 1.534 | 5.32E-07 | 0.0021 |
| 6 | **rs2858324** | 32768353 | T | 0.6397 | 7.11E-07 | 0.0016 |
| 6 | **rs2647012** | 32772436 | A | 0.6397 | 7.11E-07 | 0.0016 |

List of variants and HLA alleles in the MHC locus that are in the 99% credible set. Genome-wide significant variants, HLA alleles and amino acid residues are boldfaced.

**Figure 3.**
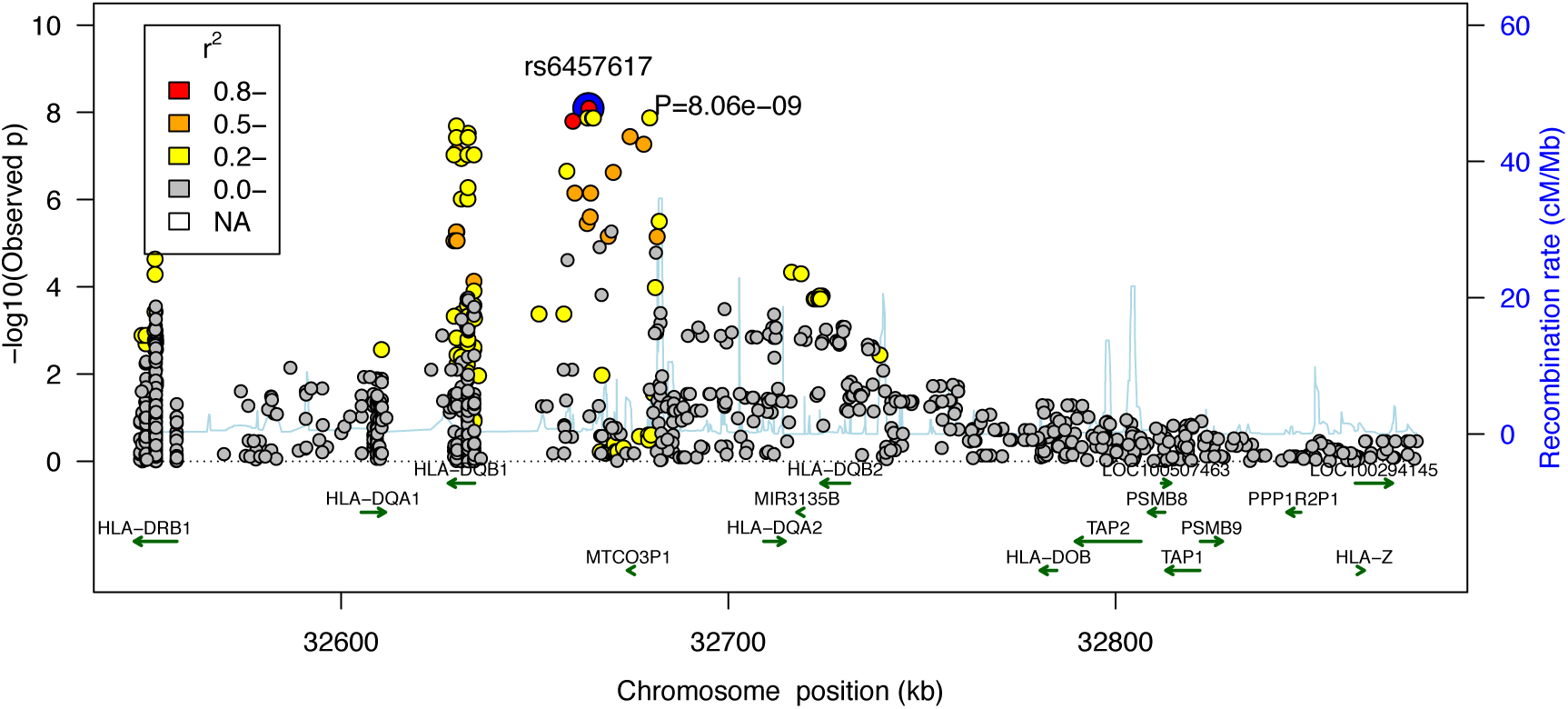
**Regional association plot for the MHC class II region.** Color indicates the linkage equilibrium with the top associated variant (rs6457620).

Neither the MHC nor the *IL28B* locus was genome-wide significant in the African ancestry. Using the heterogeneity test (fixed-effect, implemented in the R metafor package), we found that neither the MHC locus nor the *IL28B* locus have significantly different effect size (p-values =0.47 and 0.29 respectively) across the two populations. Therefore, the difference in the significance is likely driven by the sample size and/or the allele frequency differences.

In addition to the genome-wide significant loci, we examined genes outside the HLA that have been previously associated with HCV spontaneous clearance^17^. Only genes *IFNG-AS1* (p-value=6E-4) and *STAT1* (p-value=3E-4) showed marginal evidence of association (p-value < 1E-3), and this effect was observed only in the European cohort. No genes reached the marginal p-value threshold in the samples of African ancestry. *IFNG-AS1* is a long noncoding RNA that is expressed in CD4 T cells and promotes Th1 responses^18^. STAT1 is one of the key mediators of the type I, II and III interferon responses.

Since HCV is particularly diverse, with up to a 30% difference at the amino acid level between major viral genotypes, the strain of infecting virus may influence HLA-mediated clearance^13^. Unfortunately, information regarding the virus genotype or subtype was not available in this study, especially in those persons who already cleared the virus, so a direct comparison is therefore not possible. However, an indirect comparison is possible by taking advantage of the observation that North American patients are much more likely to be infected with the 1a virus and European patients are much more likely to be infected by the 1b virus{Pawlotsky:1995eg}. We observed that the association in the class II MHC locus, after accounting for the sample size difference (**Methods**), is stronger in North American samples than in European samples (Figure 4) with marginal significance (p=0.05), This suggests that viral subtype may have influenced the genetic mechanism underlying the clearance of HCV. Meta-analysis by cohorts confirms this observation (Figure 5). We also interrogated the potentially protective effect of certain SNPs associated with HLA class I alleles previously implicated in spontaneous clearance. No SNP associated with class I was associated with genome-wide significance, including those associated with *HLA B*27* subtypes (p-values > 0.05). The strength of association with the SNP most closely linked with *HLA-B*57* and control of HIV-1 (rs2395029) was not genome-wide significant but showed a marked difference by continent (North America p-value=8.6E-4, Europe p-value=0.078, overall p-value=1.0E-4), suggesting that any protective effect of this class I allele differs by region.

**Figure 4.**
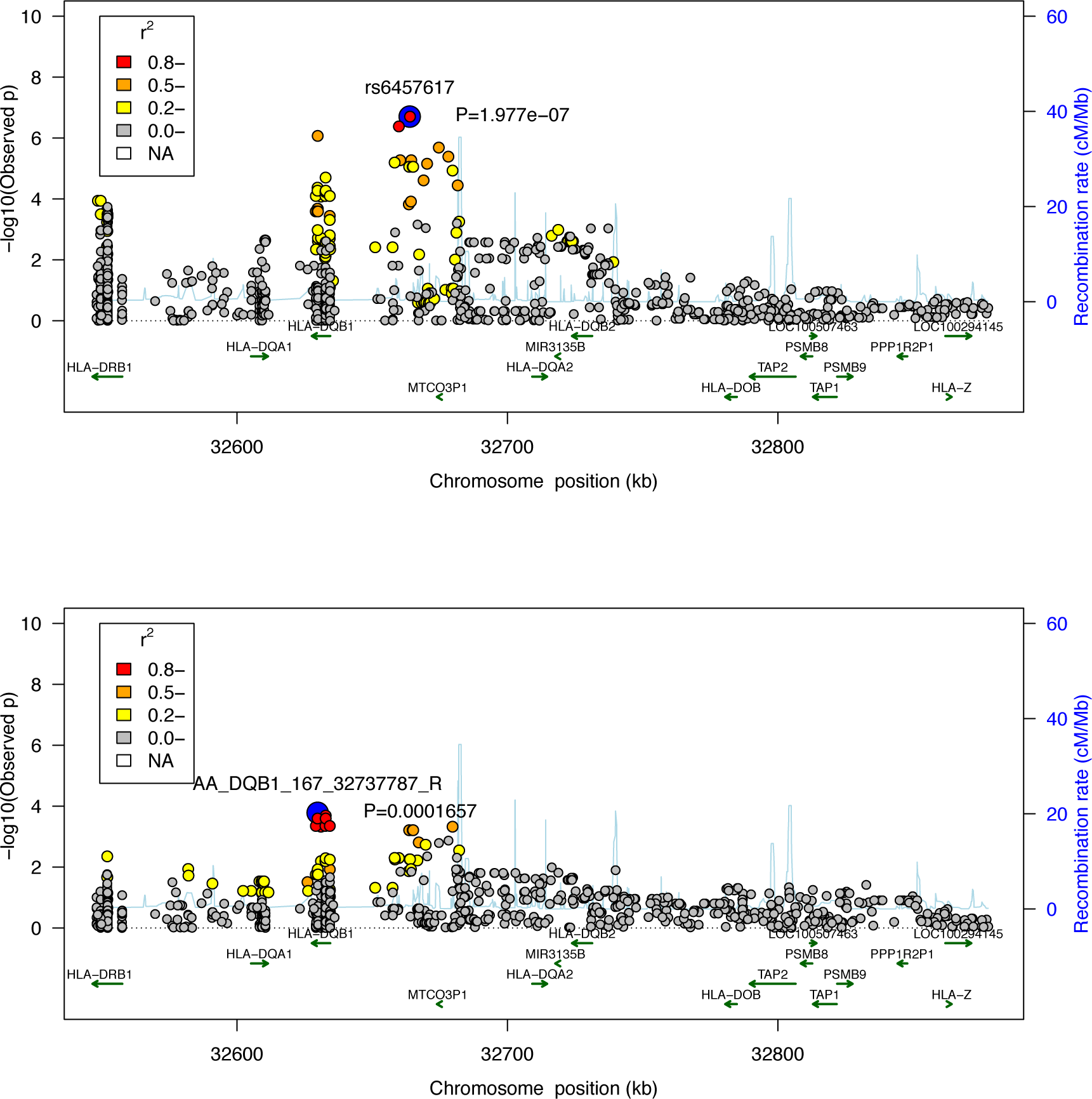
**Regional association plot for the MHC class II region in North American samples (top) and European samples (bottom).** Color indicates the linkage equilibrium with their respective top associated variant.

**Figure 5.**
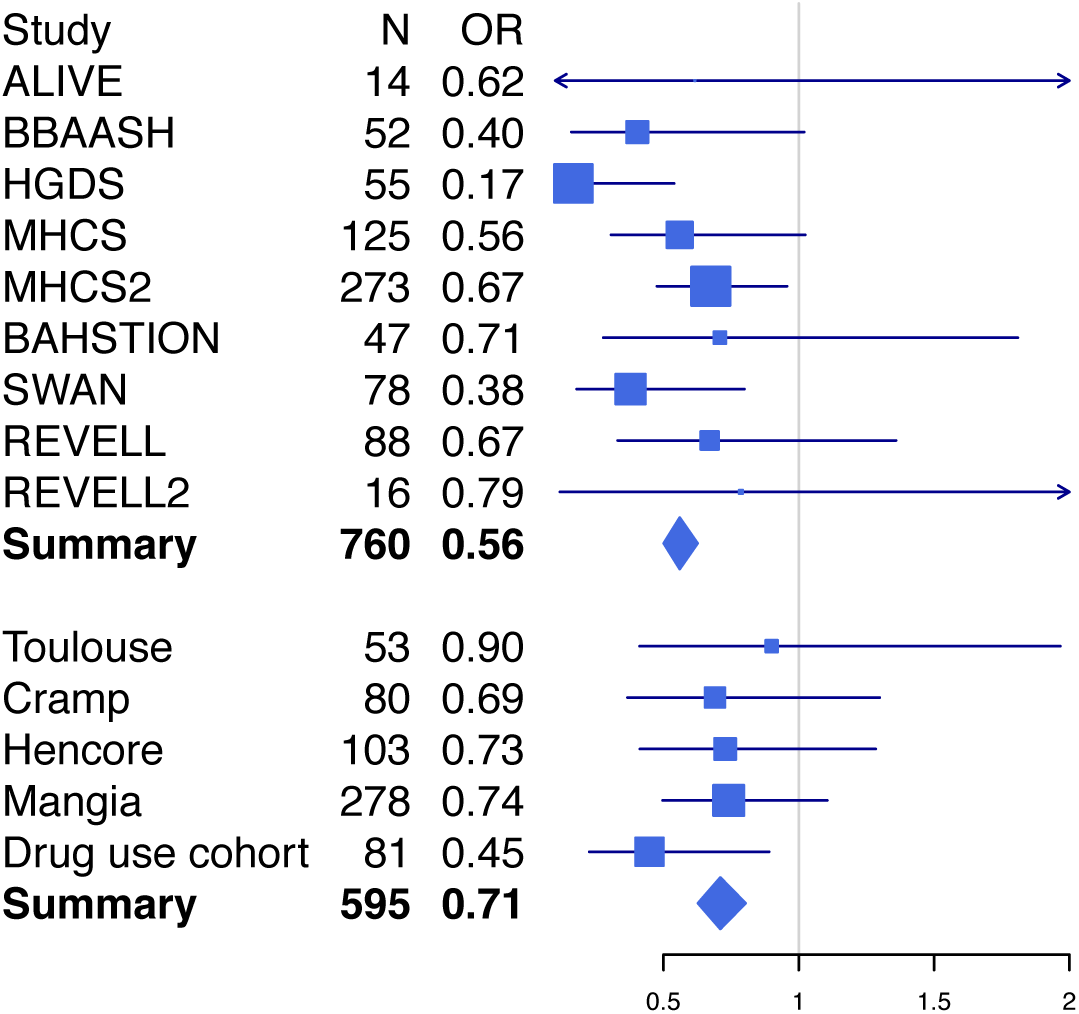
**Forest plot for the top MHC class II association (rs6457620).** Cohorts have been grouped by the geographical locations of where they were collected: the top panel includes cohorts collected in North America, and the bottom panel includes cohorts collected in Europe.

Autoimmune disorders have been reported to have shared genetic susceptibility loci^19,20^. For each of 5 major autoimmune diseases, including inflammatory bowel disease, systemic lupus erythematosus, rheumatoid arthritis, celiac disease and multiple sclerosis, we listed all variants that reached p-value<0.001 (or the best variant) in this analysis (Table 4). We found no shared variant after considering multiple testing. This analysis was only performed in the European cohort because the African cohort has limited power due to the sample size, and GWAS results in samples of African ancestry for other autoimmune disorders is more limited. A full exploration of the hypothesis that susceptibility to autoimmunity also confers ability to clear HCV will require a larger sample size.

**Table 4.** Overlap with other autoimmune disorders.

| Disease | # variant <sup>a</sup> | Source | Variant | p-value | Direction<sup>c</sup> |
| --- | --- | --- | --- | --- | --- |
| IBD<sup>28</sup> | 125/163 | Table 1 | rs2413583 | 0.0064 | + |
| SLE<sup>29</sup> | 5/6 | Table 2 | rs729302 | 0.2714 | - |
| RA<sup>30</sup> | 31/36 | Table 2 and 3 | rs11586238 | 0.0156 | + |
| Celiac<sup>31</sup> | 20/27 | Table 2 <sup>b</sup> | rs2187668 | 0.0080 | - |
| MS<sup>32</sup> | 90/97 | Table 1 and 2 | rs9967792 | 0.0159 | + |

IBD: inflammatory bowel diseases, including Crohn's disease and ulcerative colitis; SLE: systemic lupus; RA: rheumatoid arthritis; Celiac: celiac disease; MS: multiple sclerosis. <sup>a</sup>the Number of variants in this study and the number of variants reported in the publication. Not all variants are available because of QC and chip design. <sup>b</sup> Only "Previously reported risk variants" and "New loci with genome-wide significant evidence" were included. <sup>c</sup> the '+' direction means the tested allele has the same direction of effect in both diseases, and the '-' direction means the opposite.

An alternate approach, taken by the International Genetics of Ankylosing Spondylitis Consortium^21^, is to search for the reported associations with other diseases in loci having suggestive evidence (p-value < 1E-5), *i.e.*, the MHC and the *IL28B* loci in this study. We only performed the search in *IL28B* because MHC has been already implicated in many autoimmune disorders. We searched within 0.5Mb around the lead SNP (rs8099917) in *IL28B* for associations with other diseases that have been reported in the NHGRI GWAS catalog (search performed through UCSC Table Browser on August 24, 2014). Two SNPs were found to be in partial linkage disequilibrium (R^2^>0.4) with our lead SNP in *IL28B*, including rs12980275 (R^2^=0.41) associated with lipid levels in hepatitis C treatment^22^, and rs12979860 (R^2^=0.42) associated with chronic hepatitis C infection^4^/response to hepatitis C treatment^16^, which has been discussed in the previous sections.

## Discussion

We have conducted a genome-wide association study to identify genomic variants underlying the HCV spontaneous clearance using immunochip. Consistent with previous reports^4^, two loci were found to be significantly associated with the HCV spontaneous clearance in the European cohort. The immunochip design, the imputation pipeline specifically designed for the MHC region and the novel fine-mapping algorithm facilitated the accurate characterization of classical HLA types and allowed us to achieve a higher resolution in the MHC region. Twelve SNPs and 5 amino acids in the MHC region were found to be significantly associated and no secondary signal remains after conditioning on the best SNP. Fine-mapping mapped this association to a region of about 50 kilo base pairs, down from 1 mega base pairs in the previous study. We found no associated variants in the African cohort, probably due to different genetic background (in the case of the *IL28B* locus) and limited sample size (in the case of the MHC locus). Additional analyses suggested that the association in the class II MHC locus might be significantly stronger for virus subtype 1a than 1b, suggesting that viral subtype may have influenced the genetic mechanism underlying the clearance of HCV^13,23^. Additional evidence, such as virus typing, is needed to confirm this finding.

Limitations of this study include inability to dissect SNPs near the *IL28B/IFLN4* region, as this loci had not been previously implicated in autoimmune GWAS studies. While the Immunochip did include rs8099917 as a surrogate for this region, additional information regarding associations with rs12979860 and ss469415590 is not available{Kumar:2016ea}. Also, this study was a fine-mapping exercise that narrowed the MHC significantly but was not fully independent due to considerable overlap with the previous GWAS.

Previous studies of GWAS data revealed that there are SNPs and loci with evidence of association across multiple immune-mediated diseases^19^. We found several variants that have suggestive and plausible evidence of associations with both HCV spontaneous clearance and another autoimmune disorders. Despite the observation that none of these variants are significant after the strict Bonferroni correction, they jointly confirm the concept that shared genetic mechanisms underlie autoimmune disorders and suggest the hypothesis that susceptibility to autoimmunity may also confer ability to clear HCV. Fuller exploration of this hypothesis will require further analyses with larger sample sizes.

## Methods

### Overview of samples

1,944 samples from 13 cohorts (ALIVE, BBAASH, HGDS, MHCS, Rosen and colleagues, REVELL, BAHSTION, SWAN, Toulouse, Cramp and colleagues, Hencore, Mangia and colleagues, UK Drug Use Cohort) were genotyped in this study, as previously described^4^. Self-clearance of HCV was coded as cases (718 samples) and persistence of HCV was coded as controls (1,180 samples). Samples with unidentified clearance status were not used (46 samples). All samples were genotyped using Illumina’s Immunochip, a custom Infinium chip with 196,524 SNPs and small in/dels. A large number of these variants are in 187 high-density regions known to be associated with twelve autoimmune disorders and inflammatory diseases. Variants in these high-density regions include 289 established associations, variants from 1000 genome project low coverage pilot 1 study^24^, and variants discovered in re-sequencing^25^. In addition, roughly 25,000 variants were included as replication of unrelated diseases as part of the WTCCC2 project, with the purpose of serving as null SNPs in analyses.

### Sample ethnicities

To identify the sample ethnicities, we first constructed the principal component axes using Hapmap samples. 988 founders from Hapmap phase 3 (draft release 2)^26^, including samples from ethnicities ASW, CEU, CHB, CHD, GIH, JPT, LWK, MEX, MKK, TSI and YRI were used. To calculate the principal components, only common variants that are also present in the Immunochip were used, and AT/GC SNPs were excluded to avoid ambiguous strand alignment. We performed LD pruning of the variants, resulting in a total of 15,525 variants used to create the principal components. The study samples were then projected to the principal component axes and assigned the ethnicities based on their distance to the Hapmap samples. Out of 1,898 samples, 1,416 samples were mapped to European ancestry, 225 samples were mapped to African ancestry and 227 samples were admixtures and were not used in this study.

### Quality control

QC was performed separately on samples of European and African ancestries separately. Variants that failed the Hardy–Weinberg equilibrium test in controls (p-value ≤ 1E-5) or had low call rate (≤95%) were identified, and 24,820 variants were removed in European samples and 20,196 variants were removed in African samples. The remaining variants were used to perform QC in samples. Samples were cleaned for having low call rate (≤95%) or having high heterozygosity rate (> 3 standard deviations from the mean).

We then created a LD pruned dataset for calculating the identity by state (IBS) matrix and the principal components. We pruned the variants using a sliding window of 50 variants, step size of 5 variants and variance inflation factor threshold of 1.25. There were 20,782 variants in European samples, and 21,778 variants in African samples after the pruning. The IBS matrix was calculated using this LD pruned dataset and checked for sample relatedness. 28 duplicated samples in European cohorts and 9 duplicated samples in cohorts of African ancestry have been identified and removed (pi_hat>0.9). The final dataset has 527 cases and 828 controls for European cohorts, and 75 cases and 171 controls for African cohorts.

To correct for within European and within African population stratification, we calculated the principal components for samples of European ancestry and African ancestry, respectively. The first two principal components sufficiently control the population stratification in both ancestries (results not shown) and were use in the association analysis as covariates.

### Imputation

Imputation of the MHC region was performed on QC cleaned data using SNP2HLA^6^. This software package takes advantage of the long-range linkage disequilibrium between HLA loci and SNP markers across the MHC region and can perform accurate imputation of classical HLA types starting from SNP genotype data. The reference panel was created using the Type 1 Diabetes Genetics Consortium’s high quality HLA reference panel (roughly 5,000 European samples), which includes classical HLA alleles and amino acids at class I (HLA-A, -B, -C) and class II (-DPA1, -DPB1, -DQA1, -DQB1, and -DRB1) loci.

### Association test

All association tests were performed in PLINK 1.07 ^27^ using the logistic regression. We assumed additive models and used the first two principal components as covariates in the regression. HCV spontaneous clearance was coded as case so an odds ratio > 1 indicates the tested allele increases the probability of spontaneous clearance.

### Test whether an association has the same effect size across two cohorts

For a cohort of sample size *N*, assume the association of interest has p-value of *p*, and the chi-square statistic, calculated from *p* using the chi-square distribution, is χ^2^. If this association has the same effect size in the other cohort, the expected chi-square statistic in the other cohort with population *M* is approximately χ^2•^*M/N.* The significance of whether this association has the same effect size can be evaluated using the non-central chi-square distribution with one degree of freedom and non-centrality parameter of χ^2•^*M/N.*

## Acknowledgements

This project was funded in whole or in part by the National Institute of Allergy and Infectious Diseases (U19AI088791 and AI082630), the Office of AIDS Research through the Center for Inherited Diseases at Johns Hopkins University, the National Institute on Drug Abuse (R01DA033541, R01DA013324, R01DA12568, and R01DA04334). The MACS is funded by the National Institute of Allergy and Infectious Diseases, with additional supplemental funding from the National Cancer Institute (UO1-AI-35042, UL1-RR025005, UO1-AI-35043, UO1-AI-35039, UO1-AI-35040, and UO1-AI-35041). The REVELL cohort was funded by R01HL076902. Swan cohort was funded by R01-DA16159, R01-DA21550, and UL1-RR024996. The HGDS is funded by the National Institutes of Health, National Institute of Child Health and Human Development, R01-HD-41224.

## Author contributions

Study design: AYK, RTC., DLT, PD, HH; Collecting samples and clinical information: AYK, CLT, RL, JJG, AM, ALC, GDK, SM, JNB, JA, LA, SMD, MEC, SIK, LHT, MB, GJA, HRR, BRE, GML; Performed Quality Control: JA, HH; Statistical analysis: HH, PD; Writing of the manuscript: AYK, RTC, HH.

## Data availability

The data that support the findings of this study are available from the corresponding authors but restrictions apply to the availability of these data, which were used under license for the current study, and so are not publicly available. Data are however available from the authors upon reasonable request.

## Author Information

The study protocols were approved by the institutional review board (IRB) at each center involved with recruitment. Informed consent and permission to share the data were obtained from all subjects, in compliance with the guidelines specified by the recruiting center’s IRB. The authors declare no competing financial interests. Correspondence and requests for materials should be addressed to

